# Quantification of extracellular matrix components in immunolabeled tissue samples

**DOI:** 10.1101/2023.04.04.535641

**Authors:** Gerard Rubi-Sans, Marina Cler, Laura Valls-Lacalle, Agata Nyga, Soledad Pérez-Amodio, Miguel A. Mateos-Timoneda, Elisabeth Engel, Elena Rebollo

## Abstract

In recent years, the interaction between cells and the extracellular matrix (ECM) has become a new focus in understanding tissue morphogenesis, regeneration, and disease. However, the lack of specific techniques to study the ECM composition in preserved tissue structures remains a major obstacle to explaining ECM changes in response to extrinsic stimuli. To overcome this, we propose a novel strategy that uses multidimensional fluorescence microscopy and computational tools to quantify ECM composition in immunolabeled tissues and/or cell-derived matrices (CDM). This approach includes a detailed protocol for densitometric fluorescence calibration and procedures for image acquisition, processing, and automated quantification. Using this method, we present new data comparing collagen types I, III, and IV, and fibronectin contents in various tissues. These results emphasize the importance of studying ECM composition *in situ* under both normal homeostatic and disease conditions.

**Graphical abstract:** 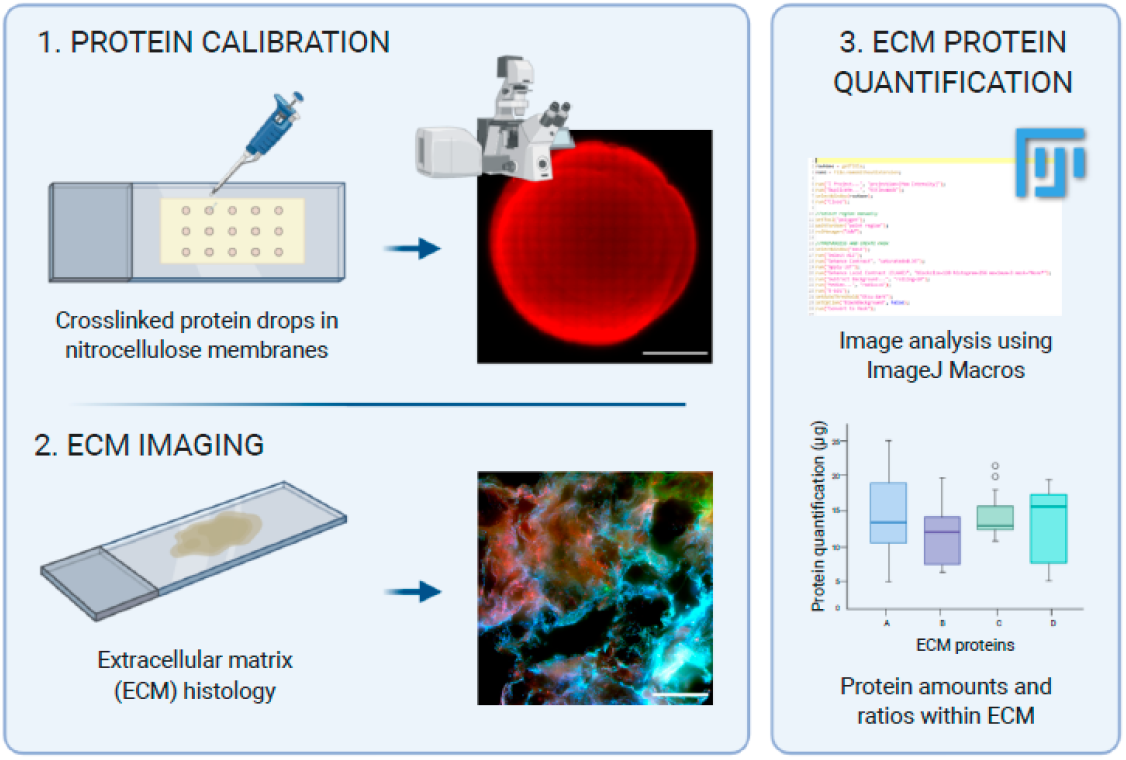

## INTRODUCTION

The extracellular matrix (ECM) is a three-dimensional macromolecular network that provides scaffolding and mediates signaling in the extracellular space to support a wide range of biological processes, such as proliferation, morphogenesis, differentiation, and homeostasis^1–3^. Understanding its composition and structure is critical in various fields, including regenerative medicine and disease diagnosis. It is a main pillar in organ engineering, which aims to develop new therapeutic approaches using biomaterial scaffolds or cell-derived matrices (CDMs). However, comparing ECM protein composition in native and decellularized tissues, as well as in artificial protein scaffolds, is challenging. The physical, topological, and biochemical composition of the ECM is not only tissue-specific but also markedly heterogeneous. Therefore, developing new approaches to study ECM composition directly in its microenvironment is imperative.

In general, biochemical methods can provide reliable information about the comparative protein contents of different tissues. For example, absorbance or fluorescence spectrophotometry can be used to measure protein concentration in bulk (Bicinchoninic acid assay or BCA) ^4^ or specific proteins such as soluble and insoluble collagens (Sircol™or hydroxyproline assays ^5,6^), elastin (Fastin Elastin assay), or glycosaminoglycans/proteoglycans (GAGs/PGs) (Blyscan or 1,9-dimethyl methylene blue (DMMB) ^7,8^ assays). Chromatography combined with mass spectrometry (MS) can fully identify the composition of tissue peptides, analyze them *in silico* and classify them ^4,9–12^. Moreover, many methods of relative and absolute MS-based protein quantification have been developed ^13^. Last, immunostaining-based techniques such as western blotting or enzyme-linked immunosorbent assay (ELISA) ^14^ have been successfully used to precisely identify and quantify ECM components. All the aforementioned techniques require sample processing that disturbs tissue architecture. Therefore, they deliver average protein amounts for the total tissue that do not shed information about its heterogeneity or protein distribution.

Advanced microscopy and image analysis provide alternative methodologies for ECM component quantification in its native environment. Histological sectioning maintains tissue structure and morphology, which enables to accurately localize and track immunolabeled ECM components without the need of disturbing their architecture. Microscopy-based approaches have been successfully used to quantify ECM components in immunohistochemistry assays using colorimetric dyes such as Picro-sirius red^15–17^, Masson’s trichrome ^18,19^, or Alcian blue ^20^. However, these dyes only distinguish between a limited number of collagen or GAGs species, and sophisticated deconvolution methods are required if individual components are to be separated ^21^. On the other hand, fluorescent or colorimetric (biotinylated or HRP-conjugated) antibodies (Abs) do allow for a much wider specific ECM components identification and provide strong potential for quantification ^22–24^. Nonetheless, there is no standardized imaging method that provides comparative quantitative data between different ECM components directly at the tissue level. We have taken advantage of the densitometric potential of fluorescence to generate a reference sample where the staining intensity linearity can be determined and correlated to protein amounts. Using this approach, we have designed a fluorescence microscopy-based protocol to quantify ECM components and compare their relative abundances using different tissues, either native or developed *in vitro*, such as CDMs ^25^, a regeneration model for skin wound healing and ischemic cardiac tissue. This protocol combines specific immunolabeling of ECM components, fluorescence densitometry, multidimensional imaging, and automated image processing and quantification. Major advances in widefield microscopy technology, both in speed and resolution, provide unprecedented grounds to study the ECM composition in the tissues of interest using this protocol.

### An overview of the method and its advantages

This method involves two main pipelines (see Figure 1): i) A densitometric fluorescence calibration step, aimed to correlate intensity with protein content (Figure 1, a-f), and ii) image acquisition and automated quantification on the tissues of interest (Figure 1, a′-d′, g, h). The calibration sample has been designed by immobilizing increasing concentrations of protein drops on top of nitrocellulose membranes. Proteins are induced to physically cross-link, generating a fibrillar mesh whose structure is comparable to that of the tissue ECM proteins. This step is key for calibration since it will enable to apply the same image processing and intensity quantification routines to both the calibration and the tissue samples. After crosslinking, protein drops are immunostained and imaged, and their total intensity is measured. For each ECM component, a linear calibration curve is built that correlates intensity with protein content. Based on this calibration, the intensity measurements obtained for each of the ECM components imaged in the biological tissues are converted into protein content. As a result, their abundances can be directly compared within the regions analyzed. A single calibration round can be used for a wide range of biological samples, given they are immunostained and acquired under the same conditions.

**Figure 1.**
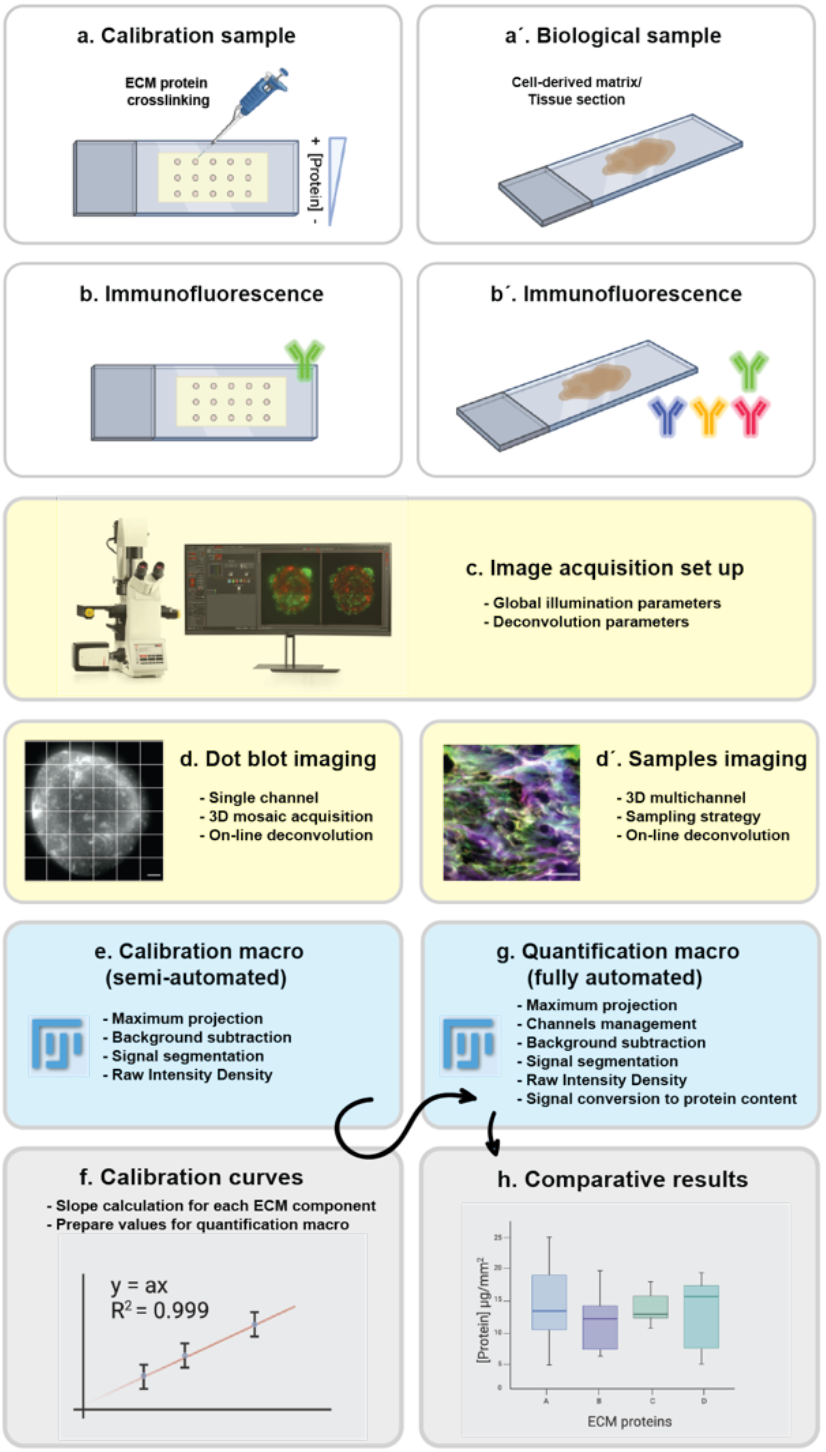
Protocol pipeline. **a**, Calibration slide, one per ECM component, containing cross-linked 0.1 μl drops of increasing concentrations of recombinant protein. **a′**, Histological tissue sections or CDMs. **b**, Each calibration slide is immunostained against its specific ECM protein. **b′**, Samples are immunostained against all studied ECM components, with the same Abs/conditions used for the individual calibration slides. **c**, A global illumination setting is configured for each fluorescent channel, to be used in both calibration and biological samples; deconvolution-assisted widefield microscopy is used. **d**, Drops are imaged using single-channel 3D mosaic acquisition. **d′**, Sample imaging is set up using multichannel 3D acquisition and a random sampling strategy. **e**, Calibration drop images are processed using a tailored Fiji macro that extracts the total fluorescence intensity per drop. **f**, Calibration curves are built for each ECM component to correlate fluorescence intensity and protein content; the calibration curves equations’ slope values are obtained to be used in the next processing step. **g**, Sample quantification is performed using a tailored Fiji macro that runs all the multichannel 3D sample images automatically and delivers the protein contents and the areas analyzed per image. An example of quantification is depicted.

Although this protocol can be performed using any microscopy instrumentation capable of optical sectioning, motorized widefield deconvolution microscopy represents the most appropriate alternative. Reduced photobleaching and high acquisition speeds are inherent features of widefield microscopy and represent a major advantage for multidimensional imaging (x, y, z, channels, tiles). Deconvolution is required to remove the out-of-focus light background generated in widefield microscopy, especially when thick samples such as tissue sections are imaged. The image contrast enhancement obtained is essential to guarantee the accuracy of measurements. When possible, GPU-assisted deconvolution should be used in parallel with image acquisition, which will save a significant amount of time. Additionally, widefield microscopes have lower costs and their maintenance routines are more affordable compared to other optical sectioning technologies such as Laser Scanning Confocal Microscopy (LSCM), Spinning Disk microscopy, or Selected Plane Illumination microscopy (SPIM). Altogether, the instrumentation chosen will speed up the imaging process and allow for the multidimensional acquisition of a high number of samples at sufficient resolution to obtain proper structure segmentation, intensity quantification, and statistics.

Last, image processing and quantification have been performed using the open-source Fiji software ^26^. We have designed two macros, for calibration and quantification respectively, using semi-automated strategies that can be easily adapted to a wide range of samples and markers. The segmentation pipeline has been designed to extract the most relevant information and can therefore be applied to a wide range of tissue structures and labels. Last, the quantification pipeline has been created for the characterization of 4 simultaneous ECM components, that can be used in any arbitrary order.

Overall, this protocol provides an accessible and efficient way to obtain protein content comparatives of several ECM components in a wide range of tissue samples. The results provided here are focused on collagen types I, III, and IV, and fibronectin, some of the most abundant and key ECM components of stromal tissues ^2^.

### Limitations

Restricted availability of commercially extracted and purified ECM proteins, PGs, or GAGs is a main constraint to study their differential expression under normal or pathological conditions. Moreover, it needs to be determined whether commercially available ECM components’ solutions can be effectively cross-linked to obtain a calibration curve. To overcome this limitation, extensive work has been performed over the years to extract and purify these high-added-value substances ^27–36^, providing researchers with efficient ways to obtain control samples for ECM components calibration.

Another limitation is the number of possible ECM components that can be compared in a single experiment. On the one hand, up to four fluorophores can be efficiently detected without signal crosstalk in filter-based epifluorescence widefield microscopes depending on the optical configuration. On the other hand, labeling up to four ECM components may require two and even three immunofluorescence staining steps, since most commercially available primary Abs are derived from mice, rabbits, and goats. These two facts will limit the quantification of a high number of ECM components in a single sample.

Less limiting but also relevant, the accuracy and reproducibility of the quantification greatly depend on the microscope setup calibration. Special attention must be paid to important parameters such as illumination stability, field homogeneity, or camera-stage alignment to avoid variations between different acquisition sessions. Additionally, a unique set of illumination parameters must be configured to acquire images of both the calibration and the biological samples. This global setting must accommodate the whole range of intensity values coming from the different samples at enough signal-to-noise ratio (SNR) to guarantee their proper quantification and comparability. Although we provide detailed recommendations on how to achieve a good general setup, poor primary Abs and labels can be a limitation in this regard.

### Experimental design

#### Calibration dot blot preparation & samples’ immunofluorescence staining

A fluorescence calibration slide must be prepared for each protein of interest, using commercially available ECM protein sources to ensure protein purity. Different concentrations (i.e., 0.1, 0.5, 0.75, 1 mg/ml) must be used to enable linear fluorescence intensity calibration. For each concentration, three to five independent 0.1 μl drops of protein solution are pipetted onto a nitrocellulose membrane. Then, proteins are cross-linked and immunostained using ECM protein-specific primary Abs (i.e., collagen types I, III, and IV, and fibronectin) and fluorescently labeled secondary Abs. Samples are then mounted for microscopy using a mounting medium and a coverslip thickness compatible with the microscope objective to be used. It is crucial to assess optimal crosslinking conditions for complete protein immobilization, determining protein absence in the crosslinking solutions’ supernatant with a standard BCA assay.

Regarding samples, the present protocol is suitable for fixed and decellularized tissues, as well as CDMs, obtained from cells cultured on biomaterial scaffolds and hydrogels ^25^. Significant differences can be found in tissue architecture and ECM morphology between decellularized and fixed native samples ^37 38^. Fixed samples are recommended when cellular and ECM morphology is also a subject of study (i.e., in tissue/organ morphogenesis studies). Otherwise, decellularized samples allow for an increased fluorescence signal and better detection of small differences between normal and diseased tissues ^39^. In any case, the procedure must be optimized for each tissue type and size, to ensure minimal alteration to the ECM structure.

Although in this protocol we are not specifically addressing sample preparation, the following key points must be considered: i) Biomaterials or hydrogels often show native autofluorescence. We suggest their removal before staining and imaging, or the use of autofluorescence reduction treatments; ^40^ ii) Thin sections (3-10μm) are recommended to improve fluorescence signal acquisition; iii) It is crucial to stain the tissue samples with the same primary and secondary Abs used in the calibration samples to obtain valid results; iv) Secondary Abs must be carefully chosen to match the available microscope filtersets configuration and avoid emission overlap; v) Primary Abs must be carefully selected to ensure specificity and avoid cross-reactivity.

#### Microscope instrumentation, setup, and image deconvolution

The present protocol can be carried out using many of the commercially available widefield microscopes having a fully motorized architecture on the main components, as named: i) xy stage, ii) z focus drive, iii) filter set configuration, iv) fast commuted light source and v) autofocus system. Such architecture allows for efficient multidimensional imaging. In the calibration samples: i) single-channel 3D mosaic acquisition (Figure 2A) is required so that the complete fluorescent drop is acquired, and its total intensity correlated to the pipetted protein content; ii) navigation interfaces able to provide overviews of the sample and/or apply automated stitching can be of great help to optimize the routine; iii) hardware and/or software autofocus tools are necessary to ensure proper focus among the tiles. Biological samples, on the contrary, can be acquired in a range of fashions depending on their structure and variability, the only requirement being that each fluorescent label must keep the same illumination conditions used during the calibration step. Therefore, a much simpler strategy can be followed using multichannel 3D stacks randomly distributed over the tissue (Figure 2B), up to a number representative enough to reflect sample heterogeneity. This will reduce file size and facilitate automation in the subsequent image quantification steps.

**Figure 2.**
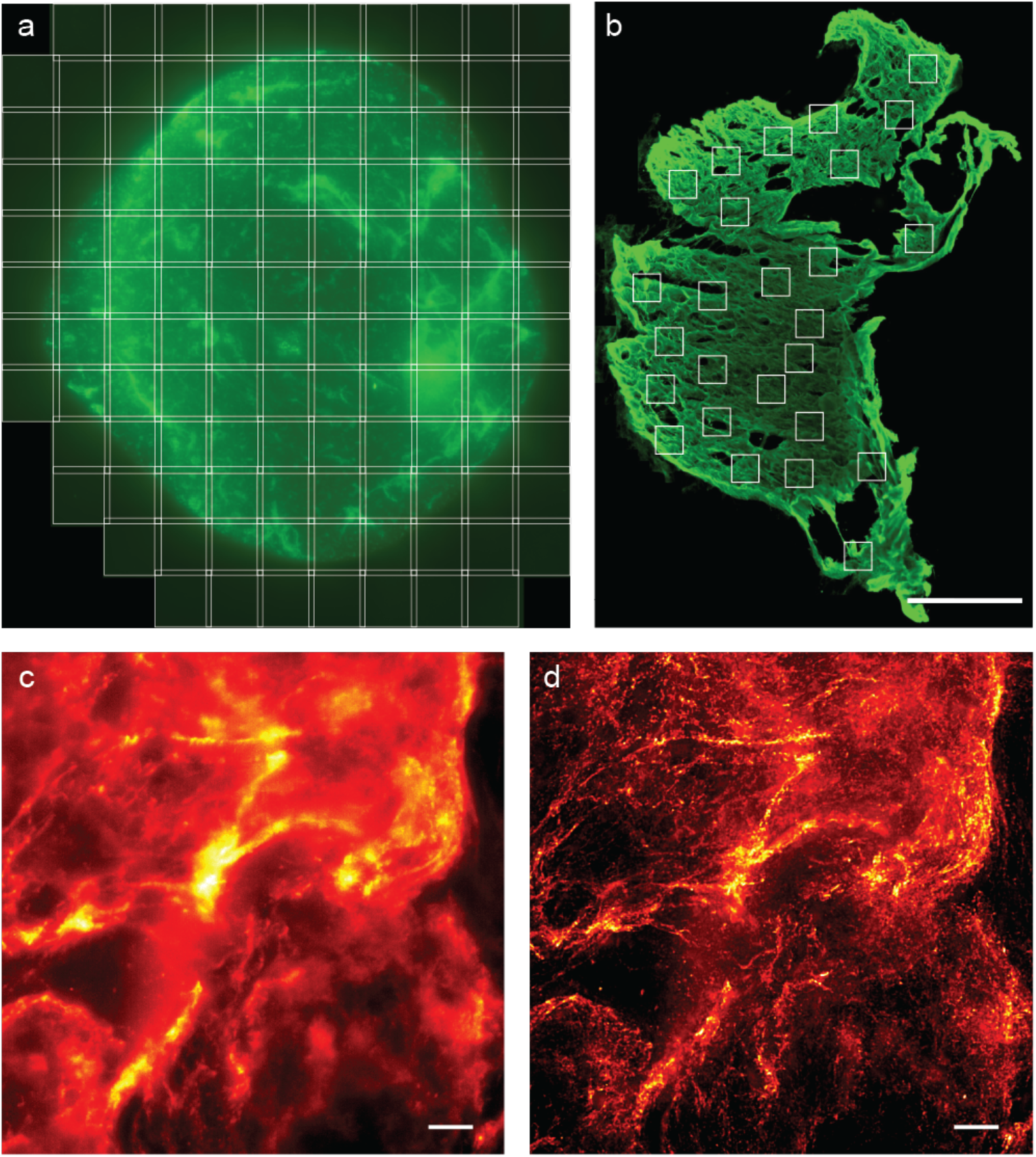
Image acquisition. **a**, Mosaic acquisition of a protein drop cross-linked onto a nitrocellulose membrane and immunostained; each square represents a 3D stack tile; each tile spans 209 μm; tiles overlap 10%. **b**, Tissue overview showing a random pattern of sampling fields (white squares) where multichannel 3D stacks will be acquired; Scale bar = 1 mm. **c**, Small area showing the 3D maximum projection of a single channel. Scale bar =10 μm. **d**, 3D maximum projection of the same region after deconvolution.

On the microscope setup, fast-changing filter configurations are required to speed up the acquisition process. Multiband dichroic mirrors combined with external fast filter wheels provide excellent speed. Additionally, barrier bandpass filters must be used to avoid emission crosstalk. Regarding detectors, modern sCMOS cameras represent a good choice due to their rapid response, high quantum efficiency, wide dynamic range, and large field of view. Choosing the right objective is critical since a high numerical aperture (NA) is needed to optically resolve small structures such as collagen fibers. To harness resolution, pixel size calibration must fulfill the Nyquist-Shannon criterium, such as the object of interest is projected on to at least 2 pixels on the camera sensor (ideally 2.5 to 3). Since image pixel size results from dividing the real pixel size in the CCD sensor by the total magnification of the system, objective magnification should be chosen accordingly. Moreover, the objective of choice should provide enough chromatic compensation for the fluorescent labels used and, ideally, coverslip thickness correction, as it is common in glycerin and water immersion lenses.

Sampling in the Z dimension must also comply with the Nyquist-Shannon criterium to guarantee the success of the subsequent deconvolution, a computationally intensive technique that will minimize the effects of the out-of-focus light generated in widefield microscopy (see Figure 2C, D). Although this is a post-acquisition step that can be performed in either ImageJ or a range of professional software (Huygens, Autoquant), GPU-assisted deconvolution in parallel to acquisition is recommended, when available, to improve the general throughput. To ensure that the quantitative data are preserved during deconvolution, iterative algorithms reassigning the out-of-focus light to their origin should be used, such as maximum likelihood estimation (MLE) algorithms ^41^. Most of the algorithms nowadays incorporated in commercial widefield microscopes are based on the regularized Richardson-Lucy algorithm ^42^, which uses the MLE mathematical strategy and is recursive and robust enough to noise.

Finally, special attention must be paid to the system calibration to guarantee the same illumination conditions over different imaging sessions. Particularly, light source stability, field illumination homogeneity, and camera-stage alignment should be kept under regular revision.

#### Image processing and quantification

Based on ImageJ/Fiji macro language we have developed two main pipelines for image processing and quantification of the deconvolved image sets. The first macro is aimed to obtain the total intensity measure from each calibration drop image. For each ECM component, a linear calibration curve must be built using the intensity measures of at least 9 drops (3 per concentration). The linear regression slopes obtained will be used to convert the corresponding intensities into protein amounts during the quantification of the biological sample images (Figure 3a-d). This is performed by the second macro, which automatically processes and quantifies all the multichannel xyz sampled images to obtain their protein contents (Figure 3e-i).

**Figure 3.**
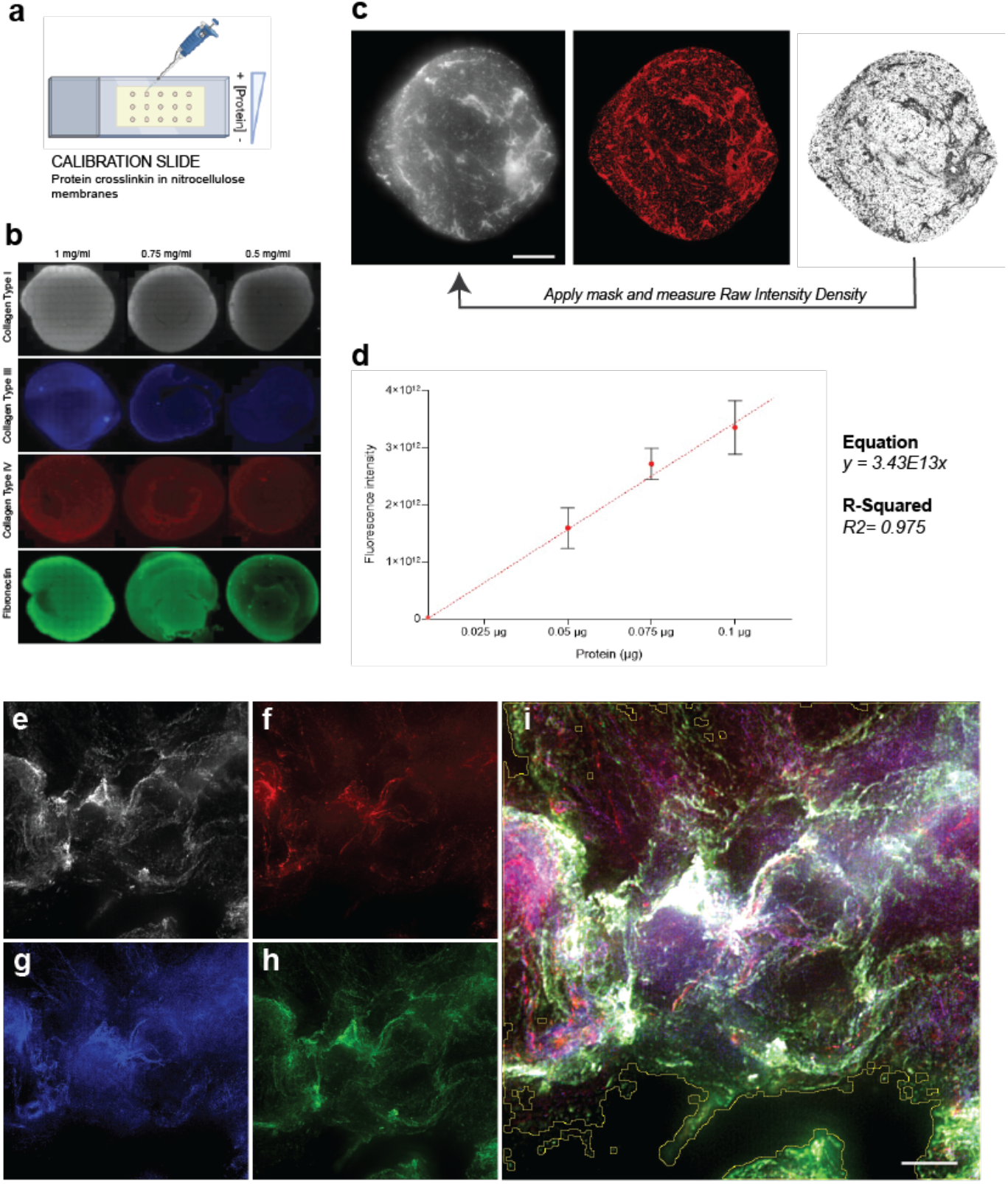
Calibration and quantification. **a**, Each calibration slide corresponds to a single ECM component and contains three to five independent drops per concentration (0.5, 0.75, and 1 mg/ml) crosslinked to a nitrocellulose membrane and immunostained. **b**, Collected images from protein drops at three different concentrations, selected from the 4 calibration slides. Scale bar = 100 μm. **c**, For each calibration drop image, the image processing macro performs the maximum projection, the background removal, and the segmentation; the resulting binary mask is used to extract the ROI to be applied on the corresponding fluorescence channel to measure the Raw Intensity Density. Scale bar = 1 mm. **d**, For each ECM component, a calibration curve is built using the intensity measurements of 3-5 independent drops per concentration; the slope of the curve will be the conversion value for the protein content quantification on the biological images. **e, f, g, h**, Maximum projection of an example image showing Collagen Type I (gray), Collagen Type III (blue), Collagen Type IV, and Fibronectin (green); Scale bar = 10 μm. **i**, After automatic segmentation a global ROI is created (yellow line) containing the most relevant tissue information; this ROI is used as a reference area for the Raw Intensity Density measurement of each channel. The obtained values are automatically converted into protein content. Normalization by area will deliver the comparative data.

The macros are available at https://github.com/MolecularImagingPlatformIBMB/TissueCollagenQuantification together with a step-by-step explanation of the image processing pipeline and a how-to-use manual. In general lines, all intensities are measured on the maximum projected images, after having automatically removed the corresponding background. Fluorescence images are further preprocessed and segmented to create the region of interest (ROIs) to be used for intensity measurements. In the biological sample images, each labeled component is separately segmented; the individual channel′s masks are then combined to create a global ROI that includes the most representative tissue region (see Figure 3i). Within the global ROI, the intensity of each ECM component is measured on its corresponding fluorescence channel. The obtained values are then converted into protein content and normalized by area for comparative purposes.

### MATERIALS

- Cell-derived extracellular matrices (CDMs) were produced in-house following the protocol described by Rubi-Sans *et al* ^25,43^
- Colorectal adenocarcinoma samples were obtained from the Hospital Clínic of Barcelona Biobank (Study approved by the Ethics Committee for Research with Drugs of the Hospital Clínic de Barcelona, Ref. HCB/2018/0174). **CRITICAL** CDMs and colorectal cancer samples (both fixed and decellularized) must be stored at 4°C in a PBS-glycine-sodium azide solution to avoid sample degradation or contamination
- Ischemic and healthy mice hearts were obtained embedded in paraffin from Vall d’Hebrón Institute of Research (Study approved by the Ethics Committee of Vall d’Hebrón Hospital and Research Institute, Ref. CEEA 35.17)
- Mice skin samples from wound healing assays were obtained at the Institute for Bioengineering of Catalonia (IBEC, Study approved by the Institutional Animal Experimentation Ethics Committee of Spain (CSIC) and Generalitat de Catalunya, Ref. CEA-OH/10727/2)
- Albumin from bovine serum (BSA, Sigma Aldrich, cat. no. A7888)
- Albumin STD Ampules (BSA STD, Thermo Scientific, cat. no. 23209)
- Ammonium hydroxide solution volumetric, 5.0 M NH_4_OH (5.0N, Sigma Aldrich, cat. no. 318612)
- Deoxyribonuclease I from bovine pancreas, lyophilized powder, Protein ≥85 % (Sigma Aldrich, cat. no. DN25)
- Ethanol absolute (VWR chemicals, cat. no. 20821.365)
- Fluoromount-G® (Southern Biotech, cat. no. 0100-01)
- Formalin solution, neutral buffered, 10% (Sigma Aldrich, cat. No. HT501128)
- Glycine, BioUltra, for molecular biology, >99.0% (NT, Sigma Aldrich, cat. no. 50046)
- Micro BCA™ Protein Assay Kit (Bicinchoninic acid, Thermo Scientific, cat. no. 23235)
- MiliQ water
- Paraformaldehyde 4% solution, EM grade (PFA, Electron Microscopy Sciences, cat. no. 157-4-100)
- PBS Tablets (Gibco-Invitrogen, cat. no. 18912014)
- Pierce™ BCA Protein Assay kit (Thermo Scientific, cat. no. 23225)
- Sodium Azide, purum p.a., >99% (Fluka, cat. no. 71290)
- Sodium phosphate 96% (Na_3_PO_4_, Sigma Aldrich, cat. no. 3424832)
- Triton® X-100 for molecular biology (Sigma Aldrich, cat. no. T8787)
- Xylene (isomeric mixture) for analysis (Merck, cat. no. 1.08297.2500)
- OptiCol™ Human Collagen Type I (Cell guidance systems, cat. no. M16S)
- OptiCol™ Human Collagen Type III (Cell guidance systems, cat. no. M20S)
- Collagen from human placenta; Bornstein and Traub Type IV (Sigma Aldrich, cat. no. C7521)
- Fibronectin human plasma (Sigma Aldrich, cat. no. F0895)
- Anti-Collagen I antibody (Mouse, Abcam, cat. no. ab6308); dil. 1:200
- Anti-Collagen III antibody (Rabbit, Abcam, cat. no. ab7778); dil. 1:200
- Anti-Collagen IV antibody (Rabbit, Abcam, cat. no. ab6586); dil. 1:200
- Anti-Fibronectin antibody (Mouse, Abcam, cat. no. ab6328); dil. 1:200
- Donkey Anti-Mouse IgG H&L (Alexa Fluor® 647, Abcam, cat. no. ab150107); dil. 1:200
- Donkey Anti-Rabbit IgG H&L (Alexa Fluor® 405, Abcam, cat. no. ab175651); dil. 1:200
- Donkey Anti-Rabbit IgG H&L (Alexa Fluor® 568, Abcam, cat. no. ab175470); dil. 1:200
- Goat Anti-Mouse IgG H&L (Alexa Fluor® 488, Abcam, cat. no. ab150117); dil. 1:500
- 2.0 mL Graduated Microcentrifuge Tubes (Neptune™, cat. no. 3765.X)
- Coverslips (No. 1.5, Duran group, cat. no. 2355035)
- Cryomolds for OCT resin
- Menzel-Gläser SUPERFROST® PLUS slides (Thermo Scientific, cat. no. J1800AMNZ)
- Nunclon™ Delta Surface, 96-well plate (Thermo Scientific, cat. no. 167008)
- Optimal cutting temperature compound (OCT, Tissue-Tek®, cat. no. 4583)
- Paraffin tissue molds
- Paraffin wax (Sigma Aldrich, cat. no. 327204)
- Spinner flask (250 mL, BellCo, cat. no. 1965-00250)

### EQUIPMENT

- Cell culture biosafety cabinet (Level 1)
- Humidified cell culture incubator at 37 °C and 5% CO_2_
- Magnetic stirrer
- Orbital shaker
- Cryostat Leica CM 1900
- Microtome Leica RM2155
- Multimode microplate reader TECAN Infinite M200 pro
- Motorized wide-field microscope Leica Thunder 3D Live Cell.
- Microscope acquisition software: LAS-X (Leica Microsystems) including online deconvolution.
- Image processing software: Fiji lifeline 22 Dec 2015 ^26^ (https://imagej.net/Fiji/Downloads) including Bioformats plugin (https://docs.openmicroscopy.org/bio-formats/5.8.2/users/imagej/installing.html)
- Fiji macros (*Protein Content Calibration*.*ijm* and *ECM Protein Content Measurement*.*ijm*) available at https://github.com/MolecularImagingPlatformIBMB/TissueCollagenQuantification
- Microsoft Excel or equivalent spreadsheet software for calculations and graphing
- GraphPad Prism 9.2.0 or equivalent statistical analysis and graphing software

### REAGENT SETUP

#### Sample decellularization solutions

Analyzed samples undergo a two-step decellularization process using a 0.1% Triton x-100 (v/v) in 25mM NH_4_OH solution followed by a 30 μg/ml DNase I solution in miliQ water. Both solutions are sterilized using a 0.2 μm filter and stored at 4 °C and -20 °C, respectively.

### EQUIPMENT SETUP

#### Microscope setup

Motorized widefield microscope Leica Thunder 3D Live cell equipped with i) HC PL APO 63x water immersion objective (NA = 1.20; WD = 0.30 mm; Coverslip correction 0.14-0.18); ii) Leica DFC9000 GTC sCMOS camera (4.2 Mpixel; 6.5 μm pixel size; QE 82%); iii) Motorized XY Stage ITK QUANTUM (Speed 500 mm/sec; resolution 5nm; Reproducibility > 1 μm; Accuracy < 1 μm); iv) SEDAT filter set configuration (XLED-QP quad filter; external excitation filter wheel with band pass filters BP440/40, BP510/40, BP 590/50 and BP 705/72); v) Spectra-X Light Engine (6 solid-state LEDs equipped with excitation filters 395/25, 440/20, 470/24, 510/25, 550/15, 575/25 and 640/30; vi) Infra-red based Adaptive focus control (AFC); vii) LasX software implemented with software navigation tools and on-line deconvolution.

#### Deconvolution setup

Although deconvolution can be performed after image acquisition, we recommend using a deconvolution routine that is fully integrated into the acquisition stream. The Leica Thunder LasX software integrates on-line deconvolution based on the regularized Richardson-Lucy algorithm and a physically modelled widefield point spread function (PSF). The algorithm has been optimized to read out local image properties during acquisition and apply the regularization based on the local environment, which corrects for further inhomogeneities in the sample while preserving the sum of all intensities ^44^. The algorithm is integrated within the so-called Small Volume Computational Clearing (SVCC) menu and can be run by simply setting to zero the computational clearing strength parameter. Fast parallel deconvolution is highly intensive and demanding in terms of hardware, a workstation is required with a minimum of 8 processors (3.6 GHz), 64 GB of RAM memory, and a graphics card similar or superior to the NVIDIA GeForce RTX-2080 Ti (11GB of video memory).

### PROCEDURE

#### Protein calibration setup and immunofluorescence staining

This subsection describes protein immobilization on nitrocellulose membranes, protein crosslinking efficiency quantification and the immunofluorescence staining procedure for calibration samples. **CRITICAL STEP** Great pipetting precision and recently calibrated pipettes are required to minimize errors and deviation in protein calibration samples.

Protein immobilization in nitrocellulose membranes. • Timing 2 hours, 40-minutes hands-on time

1. Dilute commercially available, human recombinant collagen types I, III, and IV, and fibronectin proteins to 1, 0.75, and 0.5 mg/mL concentrations using miliQ water to generate the different calibration standards.
2. Cut 1.5 × 3 cm nitrocellulose membrane pieces and place them on top of microscope slides.
3. Pipette three to five replicates of 0.1 μl from protein standards in nitrocellulose membranes for each of the tested concentrations as shown in Figure 3. **CRITICAL STEP** It is important to avoid touching nitrocellulose membranes with pipette tips, as these can be damaged.
4. Let protein drops get absorbed on nitrocellulose membranes for 15 minutes. Add 200 μl of 10× PBS to neutralize the pH and crosslink protein solutions on membranes, and induce their fibrillogenesis. Incubate samples for 1h at 37 °C.
5. Aspirate 10× PBS after protein crosslinking, and store it in microcentrifuge tubes for protein crosslinking efficiency assessment.? TROUBLESHOOTING

Bicinchoninic acid assay for protein crosslink efficiency evaluation. • Timing 1h, 20-minutes hands-on time

6. Perform a micro-BCA acid assay with 10X PBS supernatant from nitrocellulose membranes to evaluate protein crosslinking efficiency.
7. Prepare BSA standards for micro-BCA assay in 10× PBS. **CRITICAL STEP** Check manufacturer’s protocol for potential incompatible concentration. Dilute samples and/or standards if necessary.
8. Read samples’ absorbance in a spectrophotometer at a 562nm wavelength. **CRITICAL** No protein traces must be detected to ensure an optimal crosslinking on nitrocellulose membranes.

Immobilized protein dots’ immunofluorescent staining. • Timing 16 hours, 40-minutes hands-on time

9. Block immobilized protein dots on nitrocellulose membranes with a 5% BSA/0.15% glycine (w/v) solution in PBS for 30 minutes at room temperature (RT).
10. Incubate membranes with the corresponding primary Abs in a 5% BSA/0.15% glycine (w/v) solution in PBS, overnight (ON) at 4 °C.
11. Wash membranes three times, 5 minutes each, in 0.15% w/v glycine solution in PBS after primary antibody incubation. Place secondary Abs on nitrocellulose membranes in a 5% BSA/0.15% glycine (w/v) solution in PBS, at RT for 1h.
12. Wash nitrocellulose membranes three times in 0.15% glycine in PBS and microscope slides, mount slides with Fluoromount-G® mounting media, #1.5 coverslips (24×50 mm), and store them at 4 °C in absence of light until sample imaging. **CRITICAL STEP** Positive and negative controls are essential to discard protein overlapping during imaging, which would lead to incorrect fluorescence intensity results and erroneous protein quantification.

#### Sample preparation

This subsection describes fixation, decellularization and biomaterial removal for the tested samples. This protocol can be easily modified for other sample types. **CRITICAL STEP** Sterile conditions must be kept to avoid sample contamination and, therefore, potential protein quantification errors.

Cell-derived extracellular matrices (CDMs). • Timing 12 hours after CDM culture, 2-hours hands-on time

13. Obtain CDMs after a 10-day culture period, following a previously described protocol ^25^. Wash them three times for 5 minutes each in sterile PBS (no shaking) and transfer them to sterile 2 mL microcentrifuge tubes. To decellularize CDMs, add 1.5 mL of a 0.1% Triton® X-100 in 25mM NH_4_OH sterile solution, and incubate CDMs for 1h at 37 °C using orbital shaking (35 rpm).
14. Wash CDM three times in sterile PBS for 5 minutes each and perform a second decellularization step using a 30 μg/ml DNase I sterile solution for 1h at 37 °C using orbital shaking (35 rpm).
15. Remove CDM biomaterial template removal using a 250 mL spinner flask in a sterile 0.5M Na_3_PO_4_ solution for 8h at 37 °C using magnetic stirring (20 rpm).
16. Wash decellularized and biomaterial free samples three times in PBS for 5 minutes each, embed samples in OCT resin for sample histology and store them at -20 °C until further steps.

Colorectal adenocarcinoma samples. • Timing 8 hours, 1-hour hands-on time

17. Collect freshly obtained colorectal cancer samples in DMEM/F-12 HEPES cell culture medium on ice and process them within 2h in a level 1 biosafety cabinet under sterile conditions. Cut samples using surgical blades into small pieces (≈33 mm^3^) and wash them three times in sterile PBS for 2 minutes each. **CRITICAL STEP** We recommend to process tumor samples in the shortest time period possible to avoid tissue ischemia and any effect on cellular metabolism and ECM integrity.
18. Fix three to five pieces of the tumor sample in 4% PFA inside a 2 mL microcentrifuge tube for 2h at 4 °C and orbital shaking. After fixation, wash samples three times in sterile PBS for 5 minutes each, embed them in OCT resin and store them at -20 °C for histology.
19. Simultaneously, decellularize three to five tumor pieces in a 0.1% Triton® X-100 in 25mM NH_4_OH sterile solution for 4h, incubate them 1h in 30 μg/ml DNase I solution at 37 °C and under orbital shaking. Then, embed samples in OCT resin and store them at -20 °C for histology.

Ischemia reperfusion mice hearts. • Timing 34 hours after tissue collection, 1.5-hours hands-on time

20. Collect mice hearts after 14 days of reperfusion following the protocol described by Valls-Lacalle *et al* ^45^. At the end of the experiment, sacrifice animals to obtain the heart, which is sliced in 2 parts (basal region and apical region) and washed in sterile PBS to remove the remaining blood. Fix the apical region with 4% PFA for 24h at 4 °C and store it in PBS at 4 °C.
21. Dehydrate samples in a graded series of ethanol baths: one 70% ethanol bath for 1h, one 96% ethanol bath for 1h and two 100% ethanol baths for 45 minutes each, at RT.
22. Continue dehydrating tissues with two xylene baths (30 minutes each) at RT.
23. Incubate heart samples in 100% liquid paraffin two times for 1 and 4h each at 60 °C, and finally embed them in paraffin molds. Store paraffin blocks at RT for further steps.

Wound healing using poly-lactic acid (PLA) patches. • Timing 26 hours, 1.5-hours hands-on time

24. Collect mice wound healing assay tissue samples by tissue excision 14 days after generation of pressure ulcers (6 days) and wound treatment (8 days) with poly-lactic acid (PLA) or Mepilex® (Control) wound dressings, following the protocol described by Pérez-Amodio *et al* ^46^.
25. Wash excised wounds three times in sterile PBS, 5 minutes each, fix them in 10% neutral buffered formalin solution for 24h, dehydrate them in ethanol baths and embed samples in paraffin as described above.

#### Sample histology and immunofluorescence staining

OCT resin embedded biological sample. • Timing 2-hours hands-on time

26. Slice OCT-embedded CDMs and tumor samples (fixed/decellularized) at 20 μm/slice thickness using a Leica CM1900 cryostat at -20 °C.
27. Place slices on microscopy slides and store them at -20 °C until further steps.
28. Thaw OCT-embedded samples at RT and rinse them three times, 5 minutes each, in miliQ water to remove OCT resin.

Paraffin embedded biological samples. • Timing 15 hours, 3-hours hand-on time

29. Keep paraffin-embedded ischemic mice hearts and wound samples at RT before sectioning at 4 and 8 μm/slice thickness respectively, in a Leica RM2155 microtome.
30. Place slices in a 40 °C prewarmed water bath and collect them on the microscopy slides.
31. To avoid air bubbles formation underneath samples, dry samples overnight at 37 °C and store them at RT until further steps.
32. Deparaffinize samples by immersing slides twice in xylene for 5 minutes each.
33. Rehydrate samples by immersing slides in graded ethanol baths (twice in 100% ethanol and twice in 96% ethanol baths for 5 minutes each). Then, rinse samples in miliQ water until ready for staining. **CRITICAL STEP** Do not let tissue slices dry from this point on.

Sample immunofluorescence staining. • Timing 2 days, 1.5-hour hands-on time

34. Block samples after OCT/Paraffin removal for 30 minutes, using 5% BSA/0.15% glycine (w/v) solution in PBS, to avoid unspecific antibody interactions. Note that cell permeabilization is not required as only ECM proteins are being stained.
35. A two-step immunofluorescence staining takes place to stain collagen types I, III and IV and fibronectin. First, incubate samples overnight (ON) at 4 °C with collagen types I and III primary Abs in a 5% BSA/0.15% glycine (w/v) solution in PBS.
36. Wash samples three times with 0.15% glycine in PBS, 5 minutes each and incubate them 1h at RT with secondary Abs against collagen types I (647nm) and III (405 nm). Then, perform three washes of 0.15% w/v glycine in PBS, 5 minutes each.
37. Second immunostaining step starts with blocking again unspecific antibody interactions for 30 minutes, using 5% BSA/0.15% glycine (w/v) in PBS.
38. Incubate collagen type IV and fibronectin primary Abs ON at 4 °C with already stained samples. After primary Abs incubation, wash samples three times with 0.15% glycine in PBS, 5 minutes each.
39. Incubate samples with secondary Abs against collagen type IV (568 nm) and fibronectin (488 nm) in 5% BSA and 0.15% glycine (w/v) in PBS for 1h at RT and wash them again three times with 0.15% glycine (w/v) in PBS, 5 minutes each. **CRITICAL STEP** Avoid using secondary Abs with similar emission spectra to avoid fluorescence overlapping during imaging.
40. Finally, mount samples with the aqueous Fluoromount-G® mounting media and #1.5 coverslips (24×50 mm) and store them at 4 °C in absence of light until sample imaging.

#### Image acquisition

This subsection describes how to create a global illumination setup that is compatible with both the calibration and the biological samples. It is recommended to first create a multichannel configuration using the biological samples and set the optimal parameters (filter configuration, excitation power, and camera exposure time) for each of the individual fluorescent labels. The camera should be set to 16-bit when possible and image intensity should be allowed to reach an intermediate value within the dynamic range, to be able to allocate intensity variations among the different samples. Then, each individual channel should be doubled checked and further corrected using the calibration slides, to make sure that all drop concentrations have enough SNR. **CRITICAL STEP** The final channel′s configuration should be kept unchanged during both the calibration and the biological samples′ acquisition sessions.

Global illumination setup. • Timing 30-minutes to 1 hour hands-on time

41. Switch on the microscope system (on the Leica Thunder 3D Live Cell: xy stage controller, camera, light engine, microscope control box, and computer).
42. Choose a high NA objective and adjust the correction collar if needed; we have used a water immersion PLS-APO 63x NA=1.2 objective and adjusted the coverslip correction collar to 0.17 mm.
43. Focus the sample using transmission white light, i.e., bright field (BF). **CRITICAL STEP** Excessive fluorescence excitation may cause unnecessary bleaching of the fluorophores and therefore affect intensity quantification. Closing the condenser aperture may help gain some image contrast in BF imaging when no other contrast method such as polarization, phase contrast, or dark field is available.
44. Create a fluorescence multichannel configuration using as reference a biological sample having complete immunofluorescence staining. **CRITICAL STEP** It is essential to avoid signal crosstalk between channels by choosing the appropriate filter sets; acquisition speed should also be optimized; we recommend using external filter wheels containing the band pass emission filters, when available. External wheels move fast (in the order of ms) and band pass filters will allow for safe acquisition without signal contamination. **CRITICAL STEP** Signal should be adjusted within the range of linear response of the CCD sensor and below saturation level, to guarantee that intensity variations among the samples fall within the dynamic range. Since sCMOS sensors have a high dynamic range and fixed immunolabelled samples provide high signal intensities, a bit depth of 16-bit will allow working comfortably in the medium-to-high signal range while still providing a good grey level discrimination. **CRITICAL STEP** For fluorophores whose emission fall outside the chromatic correction of the lenses, relative focus correction (RFC) may be required to further correct for the chromatic mismatch.
45. Double-check the illumination settings using the calibration samples. For each channel, the corresponding calibration sample should provide images with enough SNR, especially in the low-concentration drops. Correct the excitation power and the exposure time, if necessary, to create the definitive global setting for each channel.
46. Using the calibration sample, set the 3D stack parameters; we have used 3 μm thickness and a z-slice interval of 0.26 μm. The stack dimensions must be maintained during the acquisition of both the calibration and the biological samples. **CRITICAL STEP** The distance between planes should fulfill the Nyquist-Shannon criterion allow for an efficient deconvolution.
47. Set the deconvolution parameters. In the LasX software, activate the option SVCC and set to zero the strength of the computational clearing process. The set deconvolution parameters according to figure 4. Check that deconvolution parameters work for both the calibration and the biological samples and no artifacts appear in the images.
48. Save each channel configuration separately and/or the multichannel configuration to be used for the calibration and the biological samples′ acquisitions respectively.? TROUBLESHOOTING

**Figure 4.**
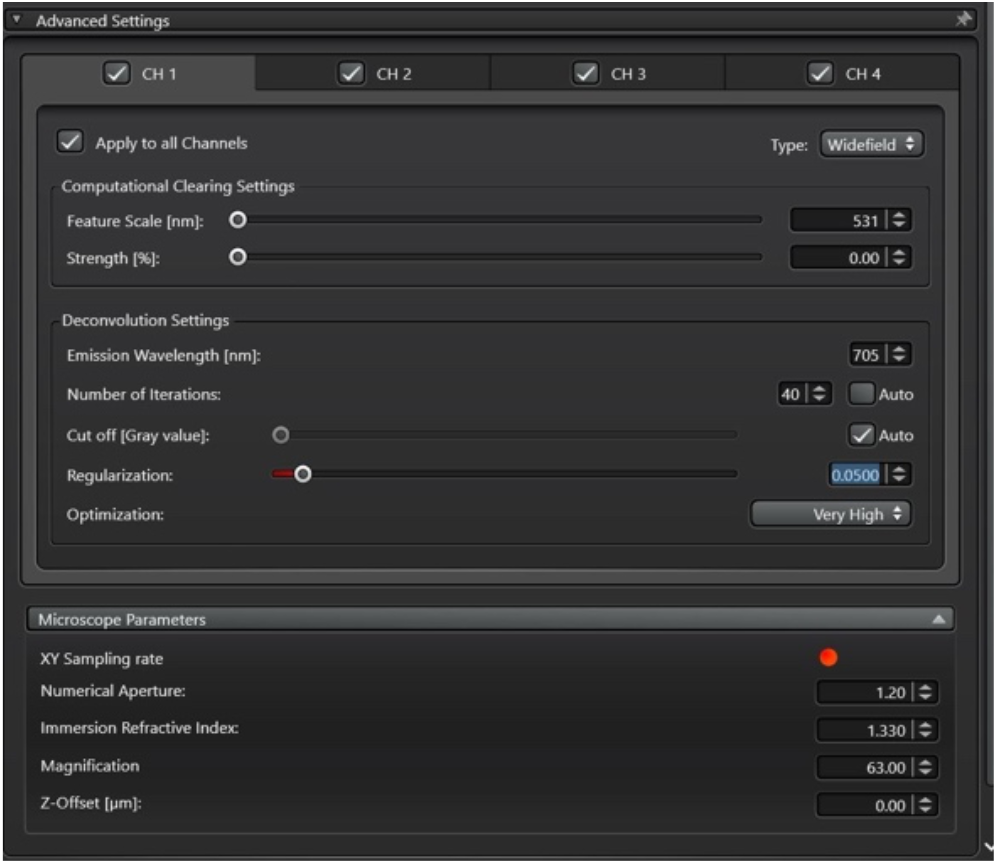
LasX deconvolution setup window. By setting the parameter “strength” to 0, the Leica algorithm “Small Volume Computational Clearing” performs a deconvolution routine based on a regularized Richardson-Lucy algorithm. The deconvolution settings can be set for each channel independently or applied to all the channels, if compatible, by clicking the option “Apply to all channels”. The number of iterations can be specified or left automatic. The cut off value subtracts the camera offset value automatically. Optimization applies a Gaussian filter to the first data point before iteration, the bigger the sigma, the higher the optimization. Finally, the optical parameters (NA, sampling rate, immersion refractive index, and distance to the coverslip are taken into account for PSF definition.

Protein calibration membranes imaging. • Timing 1-2 hours per calibration slide

49. Load the illumination configuration that corresponds to the calibration slide labeling.
50. Using a low magnification objective (i.e., 10x), acquire a preview image of the whole nitrocellulose membrane using BF to avoid bleaching. This will serve to select the region/s to be acquired as a mosaic.
51. Switch to a 63x magnification objective and focus the sample. **CRITICAL STEP** When changing from a dry to a water immersion lens make sure the sample does not move by lowering the objective and placing the water gently before focusing.
52. Adjust the mosaic acquisition conditions. This includes setting the acquisition ROIs, specifying the plane around which the z-Stack will be acquired, and checking for focus along the tiles. We recommend creating a focus map to adapt the acquisition volume to the membrane isosurface. **CRITICAL STEP** sCMOS may have a field of view bigger than the region for which the optics is corrected. We recommend using a smaller acquisition ROI (1024×1024 pixels) to avoid illumination decay towards the tiles′ edges.
53. Activate online deconvolution and automatic merging of the tiles. These two steps will considerably increase the throughput. **CRITICAL STEP** Both deconvolution and merging are postprocessing steps, although they can be considerably time-consuming and definitely a limitation if the system lacks these options.? TROUBLESHOOTING
54. Acquire 3 to 5 crosslinked protein dots per concentration tested and three or more different concentrations.? TROUBLESHOOTING
55. Save the files using the available internal format that keeps all the metadata. In the LasX software, we use *. lif extensions that can be opened right away using Fiji. **CRITICAL STEP** Since big data will have to be handled, we recommend saving the final deconvolved and merged images in separate files.
56. Repeat steps 48 to 54 for each of the proteins to calibrate.

Sample imaging. • Timing 1-2 hours per condition

57. Load the multichannel global setting
58. Choose the 63x water immersion objective and perform a single-plane overview of the tissue using one representative fluorescent channel.
59. Choose the xyz coordinates for the image sampling within the tissue on the overview. A list can be created to be done automatically. **CRITICAL STEP** Use the same acquisition ROI, z-stack, and deconvolution parameters applied to the calibration samples. **CRITICAL STEP** Choose enough fields to guarantee statistical representativity. ? TROUBLESHOOTING 1 ? TROUBLESHOOTING 2
60. Save the files using the available internal format that keeps all the metadata. In the LasX software, we use *. lif extensions that can be opened right away using Fiji. **CRITICAL STEP** Since big data will have to be handled, we recommend dividing the experiments into different files, i.e. one file per tissue, containing all the corresponding sampled stacks.
61. Repeat step 56 for 59 for the remaining tissues within the slide, and for the different conditions/slides to be imaged.

#### Image analysis and quantification

Before running the analysis, make sure the right Fiji version (Lifeline 2015 Dec 22) is installed, download the macros from the GitHub repository, and store the files in the right format. The calibration images will have to be analyzed first to obtain the conversion values. They can be kept in the internal microscope format since they will be analyzed one by one. The biological sample images will have to be converted to^*^.tif hyperstacks and stored in a reference folder for the macro to process them automatically.

Fluorescence calibration analysis. • Timing 10 minutes per image, 1 minute hands-on time, 1 hour hands-on time to build calibration curves.

62. Open one *.lif file containing a protein fluorescent dot. In the Fiji Bioformats importer choose the option to open as hyperstack. **CRITICAL STEP** Avoid autoscaling while opening to ensure the pixel values are not modified.
63. Open the macro *Protein Content Calibration*.*ijm* by dragging and dropping on the Fiji menu bar and hitting run.
64. When prompted, draw a ROI on the active image to select the reference background region.
65. When prompted, draw a polygon around the fluorescent dot.
66. Copy results to a data sheet i.e. excel.
67. Repeat steps 61 and 65 for every protein calibration dot replicate, concentration, and protein of interest.
68. Build the calibration curves plot for each protein analyzed. To that aim plot the obtained *Raw Intensity Densities* vs. the protein content and fit the results to a line that passes through zero. The slope will be used as a calibration value in the subsequent quantification step.

Protein content quantification analysis. • Timing 1-2 minutes per image

69. Open the macro named *ECM Protein Content Measurement*.*ijm* by dragging and dropping to the Fiji bar and hit *Run*)
70. A browser will pop up for the user to select the origin folder containing the images to be quantified. CRITICAL STEP They must be multidimensional images created as xycz hyperstacks in ^*^.tif format.
71. Next, a new browser will pop up for the user to select the destination folder where the results table and the verification images will be saved.
72. Then, a dialog box will show up, asking for names to identify channels C1 to C4. For each channel, the calibration curve parameters (slope and independent term, previously estimated during the calibration step) and the background level, must be introduced.
73. The macro will now sequentially process all the images from the origin folder and deliver to the destination folder a text file containing the protein concentrations per channel and per image, plus a list of verification images that can be used to track back any possible outlier result.
74. Repeat steps 69 to 72 for every folder of images to be analyzed. **CRITICAL STEP** During image analysis, compatible images (from the same biological sample) must be used as specific noise levels are subtracted.

### TROUBLESHOOTING

**Table 1.** Troubleshooting table

| Step | Problem | Possible reason | Solution |
| --- | --- | --- | --- |
| 5 | Calibration proteins cannot be crosslinked to nitrocellulose membranes | Inappropriate saline solution for protein crosslinking | Check protein manufacturer recommendations for protein crosslinking |
|  |  | Protein is unable to be crosslinked | - |
|  | Low protein crosslinking efficiency (BCA assay results show some protein in the crosslinking solution supernatant) | Not enough time for proteins to adsorb to nitrocellulose membranes | Increase the time proteins are left adsorbing to nitrocellulose membranes |
|  |  | Protein crosslinking time with saline solution is not enough | Increase protein crosslinking time with saline solution |
| 48 | Global microscope setup is not valid for calibration/biological samples | Calibration/biological samples' fluorescence intensity falls outside established microscope global setup parameters | An iterative process of setting up parameters in biological samples and test its validity for calibration samples is required for a precise and reliable protein calibration and quantification |
| 53 | Microscope lacks deconvolution tools |  | If the used microscope does not have deconvolution tools, this process must be performed after image acquisition. Professional software such as Huygens allows users to perform image deconvolution after acquisition |
| 54 | Deficient calibration protein dots' staining | Potential antibody unspecificity | Perform antibody specificity controls and test different Abs |
|  |  | Low protein signal (Microscope laser setup is not appropriate) | Once microscope lasers' configuration is set for samples of study, check if this works with calibration samples (Iterative process) |
| 59 | Biomaterial autofluorescence | Polymeric materials usually present autofluorescence | If samples of study contain autofluorescent biomaterials, consider removing the biomaterial before histology<br>Check if biomaterial autofluorescence applies for all fluorescence channels |
| 59 | Protein fluorescence signal overlapping | Two or more closely related proteins (such as collagen I and III) might lead to fluorescence overlapping | Choose primary Abs from different animal sources so the secondary Abs can attach more specifically to each protein<br>Choose secondary Abs with different excitation and emission wavelengths<br>Ensure blocking steps work properly, especially in two-step immunofluorescence staining (four or more channels)<br>Perform antibody specificity and overlapping controls before analyzing samples |

### TIMING

Steps 1-12, Protein calibration setup and immunofluorescence staining: 19 hours, 2-hours hands-on time. Steps 13-25, Sample preparation: 78 hours, 6-hours hands-on time.

Steps 26-40, Sample histology and immunofluorescence staining: 65 hours, 6.5-hours hands-on time.

Steps 41-61, Image acquisition: 1-hour hands-on time plus 2-hours hands-on time per protein calibrated and 2-hours hands-on time per sample condition analyzed.

Steps 62-74, Image analysis and quantification: 10 minutes per protein calibration image, 3 replicates per protein concentration (n=3), and 3 concentrations (total number of calibration images analyzed = 9). 1-hour to build protein calibration curves. 1-2 minutes analysis per image from analyzed samples (typically, n=10 sample regions per condition and N=3 replicates per condition).

## ANTICIPATED RESULTS

This protocol describes the procedure to quantify the amount of extracellular matrix (ECM) proteins from tissue samples and CDMs using immunofluorescence images. Crosslinked ECM protein dots (of known volume and concentration) on nitrocellulose membranes (Figure 3a) were imaged under the microscope and fluorescence intensity was quantified using an in-house developed Fiji macro. Obtained results were correlated to the amount of protein per dot to build up protein calibration curves (Figure 3b). The resulting calibration curves’ equations and their slopes (Figure 3c) were used to calculate protein amounts from fluorescence intensity values measured in biological samples of the study.

Up to this publication, we have successfully quantified collagen types I, III, and IV and fibronectin proteins from the ECM of *in vitro* developed CDMs, human colorectal tumor biopsies (both with cells and decellularized), mouse ischemic hearts, and mouse skin samples. CDM composition was assessed from three different culture strategies (Figure 5). Using this quantification method, it is possible to study differences in ECM composition due to changes in cell culture conditions, biomaterials sources, or the addition of chemicals such as macromolecular crowders (MMCs), growth factors, and other substances. Obtained results highlighted significant differences in CDMs’ collagen type I as well as in collagen type III content between tested conditions as shown by Rubi-Sans et al ^25^, suggesting that changes in CDMs’ production protocols impacted the final protein composition (Figure 5). This study used MMCs to increase ECM production by human Mesenchymal stem cells from adipose tissue (hAMSC). These MMCs showed to increase the production of collagen type I when the MMCs were added to the culture (C1 and C2 CDMs), while C3 CDM showed an increase in fibronectin that has been already correlated with the stirring conditions ^47^. Moreover, the authors performed technique validation experiments using mass spectrometry procedures to compare the obtained ratios between the four tested proteins. Quantitative mass spectrometry results showed a significant correlation with the ones obtained through protein fluorescence intensity quantification. The main advantage of the presented method is the possibility to visualize the protein distribution and colocalization. Moreover, the tailoring potential of *in vitro* ECMs, demonstrates that these can be modulated to resemble different tissues and conditions.

**Figure 5.**
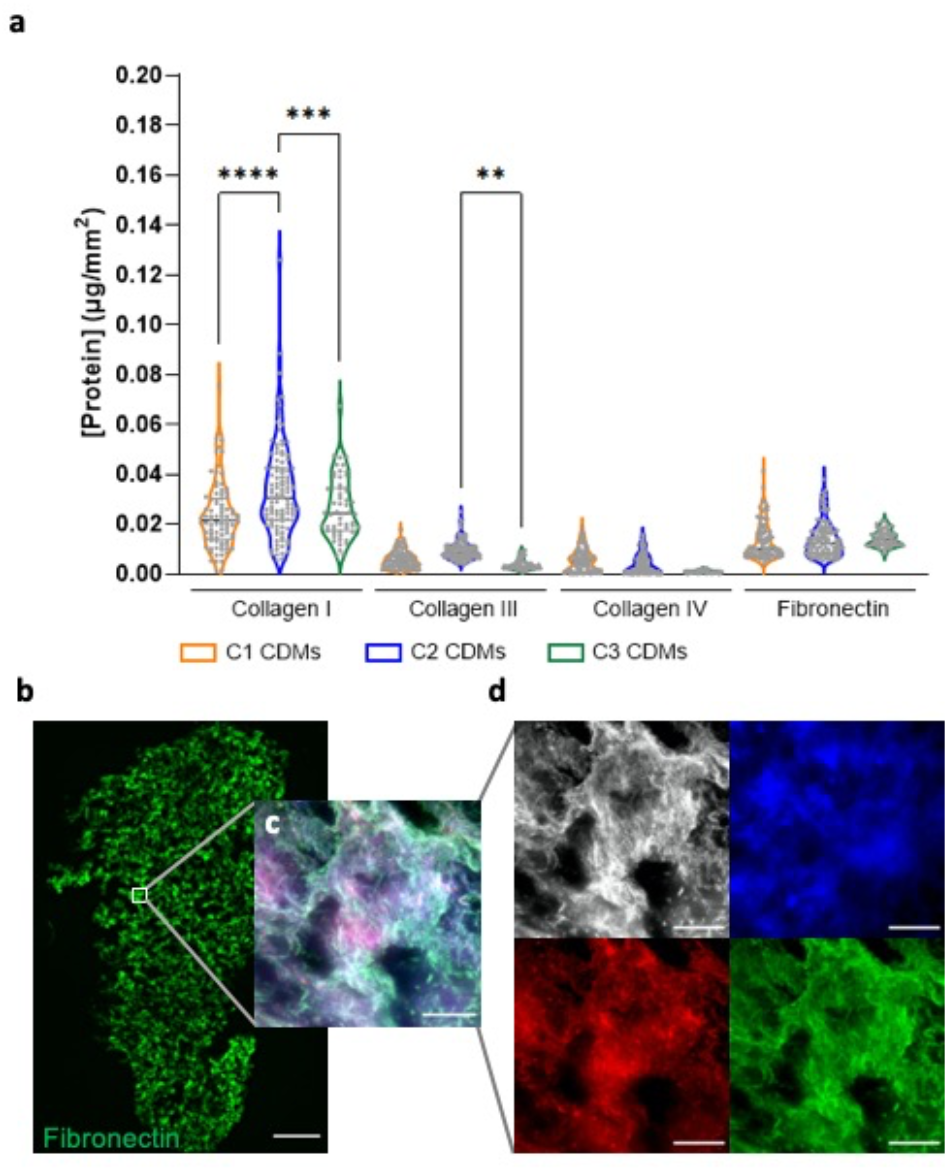
Cell-derived matrices (CDMs) protein quantification. **a**, CDM protein quantification results from three different conditions related to fabrication protocol (C1, C2, and C3). **b**, CDM tile scan of Fibronectin protein (green), scale bar = 500 μm. **c** and **d**, 3 μm Z-stack of collagen types I (grey), III (blue), IV (red) and fibronectin (green), scale bars = 100 μm. (**p-value < 0.01, * * *p-value < 0.001, * * * *p-value < 0.0001).

Next, variations in ECM composition following decellularization were studied in human colorectal cancer biopsies (Figure 6a-e). Obtained results highlighted an increase in content for each of the studied protein (collagen types I, III, IV, and fibronectin, Figure 6f) in decellularized samples. Observed changes support tissue decellularization as an effective clearing method to improve the identification and quantification of changes in the ECM through fluorescence signal ^39^ due to the significant loss of cellular protein mass ^48^. An increased interest in the tumor microenvironment (TME) has arisen in the last decade. Moreover, it has been demonstrated that besides cells, the stroma has a crucial role in tumorigenesis, cancer progression, metastasis, and therapy resistance. Stromal fibrosis and activation during cancer development result in a denser and more rigid ECM. Thus, alternative connective fibers, such as tenascin and fibronectin become more abundant, which allows cancer cells to invade through them ^49^. The identification of these specific features can help to better understand cancer disease and design more-effective anticancer therapies.

**Figure 6.**
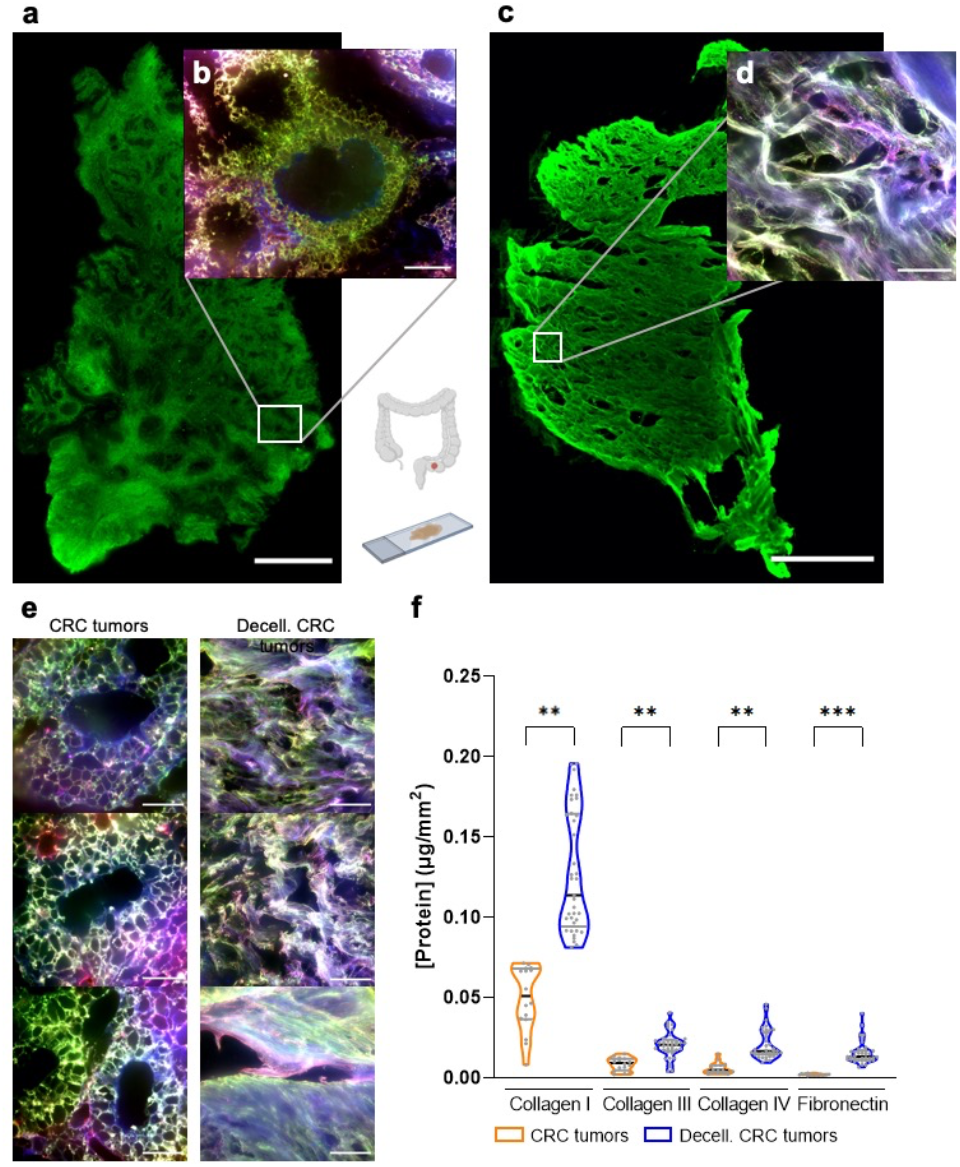
Colorectal adenocarcinoma ECM protein quantification. Tile scan of fibronectin (green) staining from **a** colorectal adenocarcinoma biopsy, scale bar = 1 mm. **b** 3 μm Z-stack images from a colorectal adenocarcinoma biopsy (collagen types I (grey), III (blue) and IV (red) and fibronectin (green)), scale bar = 100 μm. **c** Tile scan of fibronectin (green) staining from a decellularized colorectal adenocarcinoma, scale bar = 1 mm. **d** 3 μm Z-stack images from a decellularized colorectal adenocarcinoma biopsy (collagen types I (grey), III (blue) and IV (red) and fibronectin (green)), scale bar = 100 μm. **e** 3 μm Z-stack images from colorectal adenocarcinoma (Non- and decellularized) biopsies from collagen types I (grey), III (blue), IV (red), and fibronectin (green), scale bars = 100 μm. **f**, Protein quantification results. (^* *^p-value < 0.01, ^* * *^p-value < 0.001).

Furthermore, this method can also be used to study changes in the ECM during tissue morphogenesis as well as during disease development and progression (Figure 7), and *in vivo* tissue regeneration using biomaterials (Figure 8). For instance, changes in ECM composition in normal vs. ischemia-reperfusion hearts were assessed (Figure 7a-c). Results revealed no significant differences in protein composition between tested samples (Figure 7d). However, the ischemic region showed a significant loss in tissue thickness (overall mass, Figure 7b) suggesting ischemic tissue undergoes an increase in ECM fibrillar proteins density (ECM fibrosis) without a significant impact on their amount or protein abundance ratios_50_.

**Figure 7.**
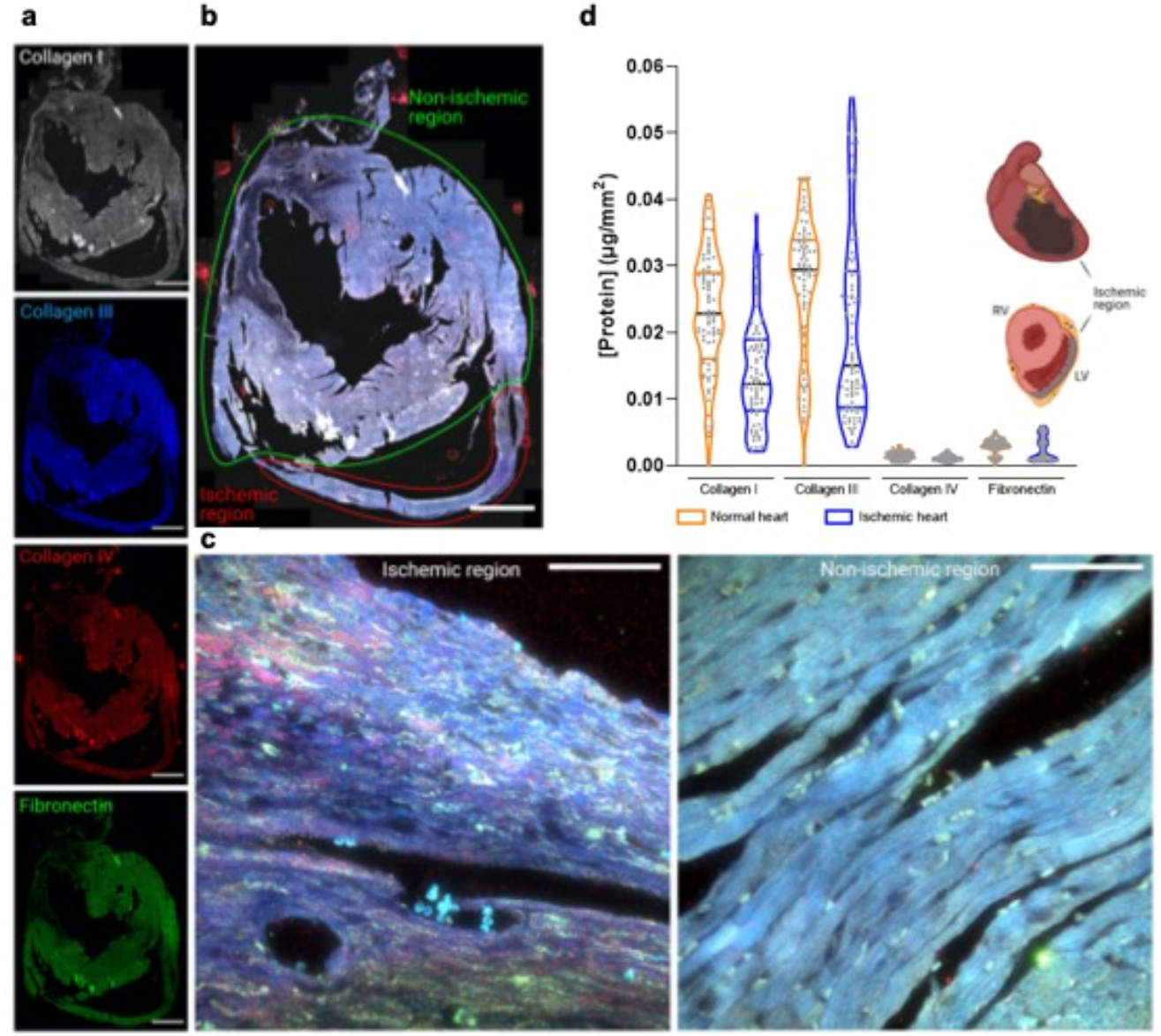
Mouse ischemic heart protein quantification. **a** and **b**, Tile scan of a collagen type I (grey), III (blue) and IV (red) and fibronectin (green) staining from a mouse ischemic heart, scale bars = 1 mm. **c**, 3 μm Z-stack from mouse’s ischemic and non-ischemic regions, scale bars = 100 μm. **d**, Protein quantification results from ischemic vs. non-ischemic regions.

**Figure 8.**
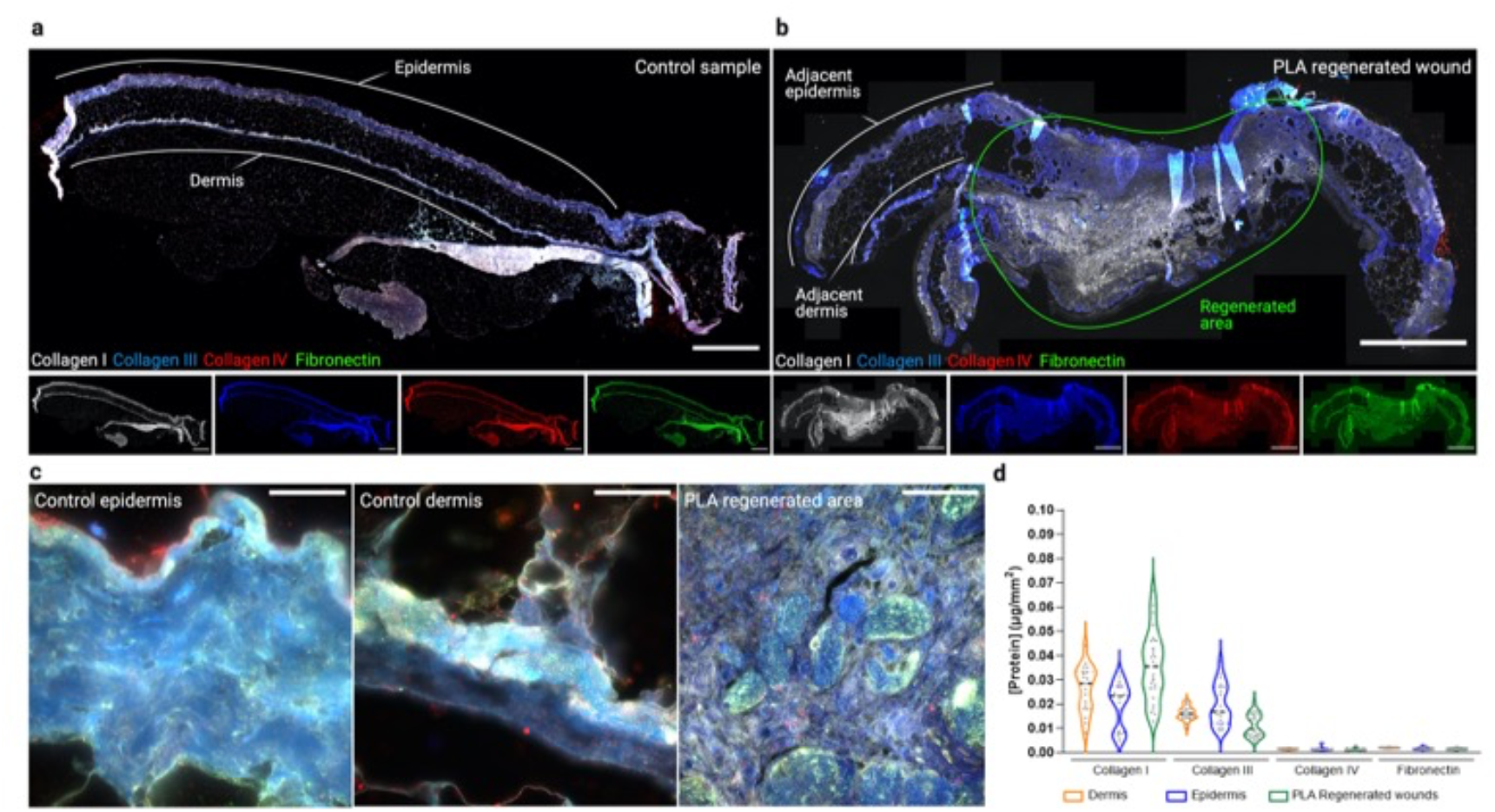
Mouse wound healing assay protein quantification. **a**, Tile scan of a collagen type I (grey), III (blue) and IV (red) and fibronectin (green) staining from a control sample mouse wound healing assay and **b**, from a PLA wound dressing regenerative approach, scale bars = 1 mm. **c**, 3 μm Z-stack from control mouse’s epidermis and dermis areas, and regenerated areas, scale bars = 100 μm. **d**, Protein quantification results from control vs. PLA wound dressing approaches for wound regeneration.

In an example of *in vivo* tissue regeneration process, a poly-lactic acid (PLA) wound dressing was used to regenerate mice skin (Figure 8). An untreated control sample (Figure 8a,c) was compared to a PLA regenerated wound (Figure 8b,c). Obtained results highlight no significant differences in ECM protein content in PLA regenerated wounds compared to control dermis and epidermis tissues (Figure 8d), suggesting PLA patches effectively promote skin ECM deposition of similar composition to that of control samples. Nevertheless, important morphological differences were observed between control and PLA regenerated samples, with the latter ones displaying no stratification of skin structures (dermis and epidermis), due to a rapid regeneration process involving the formation of granulated tissue in the wound healing process. Therefore, the use of this method not only demonstrates a correct wound healing process through protein quantification but it also can be used to follow up with the different phases of tissue healing.

When ECM components of interest are successfully calibrated, imaged, and quantified, this protocol provides researchers with valuable information about ECM composition and protein abundance from *in vitro* and/or *in vivo* experiments. The study of ECM changes over time in tissue samples and *in vitro* models is crucial in understanding disease development and progression, such as during cancer, neurodegenerative, autoimmune, cardiovascular, or connective tissue diseases. Correlation between the quantification of the ECM changes and protein localization in the tissue, with molecular and mechanical alterations in cell and tissue properties could provide a major breakthrough in identifying new therapeutic targets and developing more specific patient treatments.

## Reporting summary

Further information on research design is available in the Nature Research Reporting Summary linked to this article.

## Data availability

Raw data supporting this study is available from the corresponding authors upon reasonable request.

## Code availability

Protein calibration and protein quantification macros can be found at https://github.com/MolecularImagingPlatformIBMB/TissueCollagenQuantification

## Acknowledgments

Researchers acknowledge Dr. Antonio Rodríguez Sinovas from the Cardiovascular diseases group at Vall d’Hebrón Institute of Research (VHIR) for providing mice heart histological samples. Imaging has been performed at the Molecular Imaging Platform IBMB (CSIC)-PCB, in a Leica Thunder 3D Live Cell supported by CSIC (Bolsa de Apoyo Excepcional 1501/18) and Leica Microsystems.

## Author contributions

Conceptualization and Methodology: G.R.S., E.E., and E.R. Software: E.R. Formal analysis: G.R.S. and ER. Investigation: G.R.S. M.M.H., L.V.L., A.N., M.A.M.T. and ER. Writing - Original Draft: G.R.S. and E.R. Writing-Review & Editing: G.R.S., E.E., and E.R. Funding acquisition: E.E. and E.R.

